# The first two chromosome-scale genome assemblies of American hazelnut enable comparative genomic analysis of the genus *Corylus*

**DOI:** 10.1101/2023.04.27.537858

**Authors:** Scott H. Brainard, Dean M. Sanders, Tomas Bruna, Shengqiang Shu, Julie C. Dawson

## Abstract

The native, perennial shrub American hazelnut (*Corylus americana*) is cultivated in the Midwestern U.S. for its significant ecological benefits, as well as its high-value nut crop. Implementation of modern breeding methods and quantitative genetic analyses of *C. americana* requires high-quality reference genomes, a resource that is currently lacking. We therefore developed the first chromosome-scale assemblies for this species using the accessions ‘Rush’ and ‘Winkler’. Genomes were assembled using HiFi PacBio reads and Arima Hi-C data, and Oxford Nanopore reads and a high-density genetic map were used to perform error correction. N50 scores are 31.9 Mb and 35.3 Mb, with 90.2% and 97.1% of the total genome assembled into the 11 pseudomolecules, for ‘Rush’ and ‘Winkler’, respectively. Gene prediction was performed using custom RNAseq libraries and protein homology data. ‘Rush’ has a BUSCO score of 99.0 for its assembly and 99.0 for its annotation, while ‘Winkler’ had corresponding scores of 96.9 and 96.5, indicating high-quality assemblies. These two independent assemblies enable unbiased assessment of structural variation within *C. americana*, as well as patterns of syntenic relationships across the *Corylus* genus. Furthermore, we identified high-density SNP marker sets from genotyping-by-sequencing data using 1,343 *C. americana, C. avellana*, and *C. americana* x *C. avellana* hybrids, in order to assess population structure in natural and breeding populations. Finally, the transcriptomes of these assemblies, as well as several other recently published *Corylus* genomes, were utilized to perform phylogenetic analysis of sporophytic self-incompatibility (SSI) in hazelnut, providing evidence of unique molecular pathways governing self-incompatibility in *Corylus*.

## Introduction

Hazelnut (*Corylus* spp.) is a globally significant nut crop, with annual production of over 1 million tons, produced across 34 countries. This production is driven by a wide range of market uses across diverse industries, including food products, pharmaceuticals, and dietary supplements (Ceylan et al., 2022). Currently, commercial cultivation relies overwhelmingly on cultivars of European hazelnut (*C. avellana*) adapted to Mediterranean climates (Di Lena et al., 2022) with moderate winters to satisfy their chilling requirements (Mehlenbacher, 1991). The distribution of hazelnut production reflects these climatic requirements, with major centers of production in Turkey, Italy, and Spain, with Turkey alone producing over 65% of the global supply (Semih Uzundumlu et al., 2022). In the U.S., this narrow range of adaptation currently restricts production to relatively small regions such as the Willamette Valley in Oregon (Revord et al., 2020).

As a woody perennial crop, hazelnuts have substantial potential to provide ecosystem services such as reduced soil erosion and nutrient runoff (Demchik et al., 2014), while also sequestering carbon in above and belowground biomass (Granata et al., 2020). Expanding the climatic conditions under which commercially viable hazelnut varieties can be grown would not only make production more resilient in the face of changing climates, but provide high-value alternative crops to producers across wider geographies. One possible method for the development of commercially viable hazelnut varieties for additional growing regions, is the improvement of the native North American shrub *C. americana*, which possesses disease resistance and cold hardiness traits lacking in most cultivated forms of European hazelnut (Molnar et al., 2018). To this end, genomics-aided breeding of *C. americana*, utilizing methods such as marker-assisted selection and genomic prediction, would be greatly aided by a high-quality reference genome assembly. Such a resource would, for example, facilitate the rapid identification and alignment of genetic polymorphisms across experimental and breeding populations (Kang et al., 2016). In addition, well-annotated assemblies provide the opportunity to investigate the genetic components underlying specific biomolecular pathways involved in abiotic and biotic stress resistance, employing methods ranging from fine mapping of quantitative trait loci (QTL) to gene editing (Bolger et al., 2017; Dmitriev et al., 2022).

To facilitate such applied and experimental approaches, this study reports the first chromosome-scale reference assemblies for *C. americana* using two accessions, ‘Rush’ and ‘Winkler’, which have historically been widely used in hazelnut breeding in the Eastern United States, largely as sources of resistance to the endemic fungal pathogen *Anisogramma anomala* (Eastern Filbert Blight; EFB) (Bhattarai et al., 2017). Constructed using a combination of long-read sequencing, chromosome conformation capture, and genetic mapping, these chromosome-scale reference assemblies are of high-quality and can immediately be deployed in advancing breeding and genetic research objectives. Such potential is illustrated in this study through the identification of high-density single nucleotide polymorphism (SNP) markers for over 1,343 hazelnut plants, drawn from Midwestern natural and breeding populations. These markers provide a detailed assessment of population structure in *C. americana* and interspecific hybrids between *C. americana* and *C. avellana*. 20^th^ century hazelnut breeding the Midwestern U.S. involved interspecific hybridization with *C. avellana* (Weschcke, 1954; Rutter, 1987), but generations of open-pollination and lack of detailed pedigree records obfuscate the genetic background of current Midwestern varieties. An analysis of population structure helps to resolve this question. Understanding the proportional representation of *C. americana* and *C. avellana* genetic backgrounds in varieties which have been successful selected under Midwestern conditions will help determine breeding strategies for key traits.

In addition, the last two years has seen the release of chromosome-scale genome assemblies for several other *Corylus* species: *C. avellana* (cultivars ‘Tombul’ (Lucas et al., 2021) and ‘Tonda Gentile della Langhe’ (‘TGdL’) (Pavese et al., 2021)), *C. heterophylla* (accessions from Siping City, Jilin (Liu et al., 2021) and Yanqing, Beijing (Zhao et al., 2021)), and a wild specimen of *C. mandshurica* (Li et al., 2021). Together with the two *C. americana* genomes reported here, this study presents the first comparative genomic analyses across nearly half of the *Corylus* genus.

Comparative genetic analysis also is useful for analyzing specific traits, such as self-incompatibility, an understanding of which is critical to both hazelnut breeding and production. Plants have evolved numerous strategies to increase the frequency of outcrossing, ranging from variation in the timing of floral development, to genetically regulated mechanisms. With respect to the latter, two primary classes of self-incompatibility have been identified: gametophytic self-incompatibility (GSI) and sporophytic self-incompatibility (SSI) (Silva & Goring, 2001). The former is typified by *Rosacea* and *Solanaceae* species, wherein the haploid genotype of the male gamete at an “S-locus” (often encoding an F-box (SLF/SFB) protein) is detected by the female parent, which blocks the growth of the germinating pollen tube (Sijacic et al., 2004; Sassa, 2016). The latter is exemplified by *Brassicaceae* species, where the diploid genotype of the male parent at the S-locus (often encoding a cysteine-rich protein (SCR/SP11)) mediates detection by the female parent, and a suppression of pollen germination (Hiscock & McInnis, 2003). Self-incompatibility in *Corylus* is a form of SSI, regulated by a single S-locus (Mehlenbacher & Thompson, 1988), which fluorescent microscopic observation of pollen tube growth has been demonstrated to include at least 33 alleles with 8 levels of linear dominance (Mehlenbacher, 1997, 2014). We pursue a phylogenetic approach to the analysis of SSI using the transcriptomes reported here, and this analysis sheds additional light on the evolution of this mechanism in *Corylus*.

## Results

### Sequencing and assembly

Table 1 summarizes the raw sequence data used in assembling the two genomes. For ‘Rush’, 14.97 Gb of CCS sequence (44.32x coverage) and 13.60 Gb of ONT sequence (40.29x coverage) were generated. For ‘Winkler’, 19.60 Gb of CCS sequence (58.02x) and 12.69 Gb of ONT sequence (37.57x coverage) was generated. Short read sequencing of Arima Hi-C libraries generated 78.7x and 114x coverage for ‘Rush’ and ‘Winkler’, respectively. While ‘Winkler’ libraries therefore generated substantially more data across all sequencing platforms, in the case of both accessions, coverage was judged to be more than sufficient for the assembly of a diploid organism, assumed to have a relatively small genome size of ∼350-370 Mb, based on comparisons with previous *Corylus* assemblies (Li et al., 2021; Liu et al., 2021; Lucas et al., 2021; Pavese et al., 2021; Zhao et al., 2021). The contact map for ‘Winkler’, showing contigs scaffolded into chromosome-scale pseudomolecules, is shown in Fig. 1C (visualized using Juicebox (https://github.com/aidenlab/Juicebox)).

**Figure 1.**
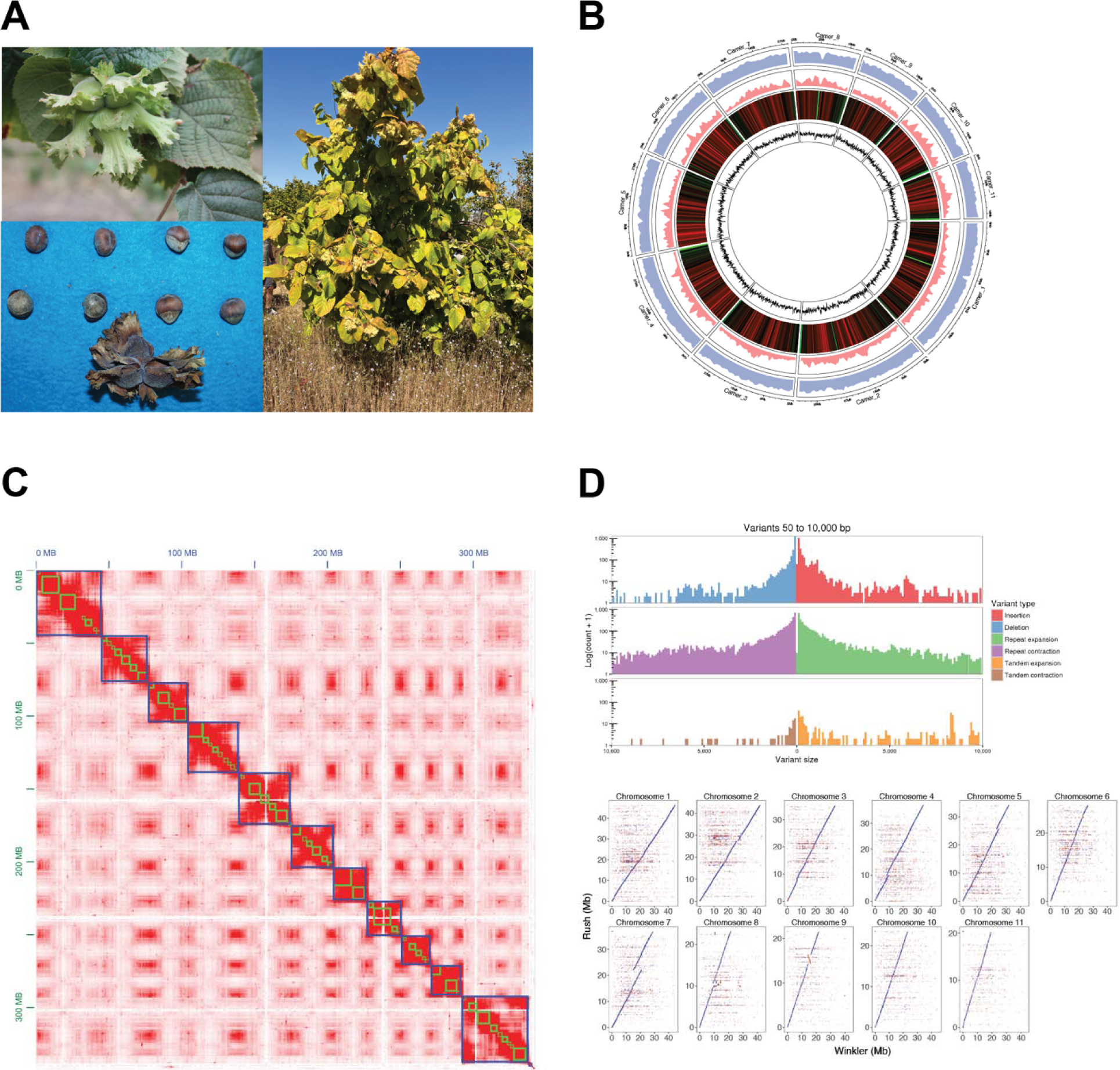
**A:** Clockwise from top left: the typical involucres, growth habit, and nuts of *C. americana* ‘Rush’, held by the NCGR in Corvallis, Oregon; **B:** Circular plot of genomic features of ‘Winkler’. From outside in: ideogram of the 11 chromosomes of *C. americana*; total gene content, as identified by gene prediction and annotation; transposable elements, as identified by RepeatModeler; heatmap of tandem repeats; line graph of tRNA content. **C:** Contact map showing aligned and sorted Arima Hi-C data for ‘Winkler’ as visualized in Juicebox; green squares indicate ordered and oriented contigs, and blue squares indicate chromosomes. **D:** Whole genome comparison of the ‘Rush’ and ‘Winkler’ chromosome-scale assemblies. Upper plot: Histograms of insertion/deletions, repeat expansions/contractions, and tandem repeat/expansions as visualized by Assemblytics; lower plot: Dot plot of delta file generated by nucmer. While small indels are evident, largely consistent synteny is observed across all 11 chromosomes. A single sizeable inversion on chromosome 9 is highlighted in red.

**Table 1.**
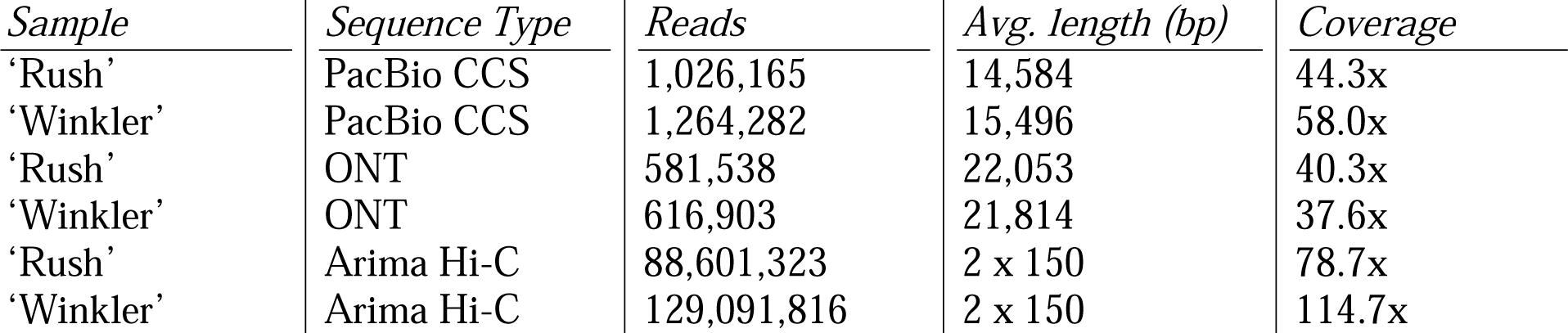
Summary of sequence data generated on PacBio, Oxford Nanopore and Illumina sequencing platforms.

Table 2 presents statistics summarizing the quality and contiguity of the ‘Rush’ and ‘Winkler’ assemblies. While both genomes exhibit high quality and contiguity, with N50 scores > 30 Mb, and over 90% of the genome assembled into 11 pseudomolecules, for both accessions there are clear differences which reflect the divergence in assembly methods. Both assemblies are close in length to previous reports of the size of *Corylus* species, however the ‘Rush’ assembly is over 50 Mb larger than the ‘Winkler’ assembly. This is likely due to unresolved duplication, apparent also in Fig. 1D, due to the fact that Hi-C sequence data could not be included in the assembly, which is also reflected in the many more contigs present in ‘Rush’ compared to ‘Winkler’. While including Hi-C data for ‘Winkler’ led to a reduction in total contigs from 398 (following hifiasm and haplotig_purge) to the 264 reported in Table 1, including Hi-C data for ‘Rush’ led to an expansion and fragmentation of contigs to 1,845. Benchmarking Universal Single Copy Orthologs (BUSCOs) were calculated for ‘Rush’ and ‘Winkler’, and while both assemblies had nearly identical complete and single copy BUSCOs represented (95.1% and 94.6%, respectively) ‘Rush’ contained more duplicates (3.8% vs. 1.4%). This finding, and the slightly larger maximum contig size also suggest unresolved artificial duplication.

**Table 2.** Summary of statistics related to genome assembly. BUSCO score codes: C: complete; S: complete and single copy; D: complete and duplicated; F: fragmented; M: missing.<colcnt=1>

| *Statistic* | *‘Winkler’* | *‘Rush’* |
| --- | --- | --- |
| Assembly size (bp) | 337,645,099 | 388,211,906 |
| Number of contigs | 264 | 1118 |
| GC content (%) | 36.05 | 36.68 |
| Assembly L50 (#) | 5 | 5 |
| Contig N50 (bp) | 31,913,876 | 35,381,848 |
| Maximum contig size (bp) | 44,790,255 | 46,319,783 |
| Genome in chromosomes (%) | 97.1 | 90.2 |
| BUSCO (C [S, D]; F; M) (%) | 96 [94.6, 1.4]; 0.9; 3.1 | 98.9 [95.1, 3.8]; 0.2; 0.9 |

Both assemblies were also annotated for several genomic features: gene models were predicted using a combination of RNAseq data and protein homology data; transposable elements were identified using RepeatModeler; tandem repeats were identified using TandemRepeatFinder; and tRNAs were predicted using tRNAscan-SE. These features are presented in a Circos-style plot in Fig. 1B. Specific statistics related to the gene model predictions are summarized in Table 3.

**Table 3.** Summary of statistics for the annotation of ‘Rush’ and ‘Winkler’.

| *Statistic* | *‘Winkler’* | *‘Rush’* |
| --- | --- | --- |
| Primary transcripts | 23,331 | 24,562 |
| Alternative transcripts | 5,701 | 6,174 |
| Median exon length (bp) | 159 | 159 |
| Median intron length (bp) | 245 | 248 |
| Average number of exons per gene | 5.7 | 5.7 |

These annotations appear extremely similar in terms of the structure of the gene models which were predicted. Slightly fewer genes were predicted in ‘Winkler’, which in this case is likely not the consequence of unresolved duplication, as the genomes were soft masked for repetitive elements. Finally, additional statistics related to the genome annotation are provided in Table 4, which compares the ‘Rush’ and ‘Winkler’ assemblies to the five currently published chromosome-scale *Corylus* assemblies.

**Table 4.** Comparison with annotations of previously published *Corylus* species. BUSCO scores refer to all complete representatives, both single and duplicated.

| *Species and cultivar name or accession* | *Genome size (11 chromosomes)* | *Repeat content* | *Predicted genes* | *Genome BUSCO* | *Annotation BUSCO* |
| --- | --- | --- | --- | --- | --- |
| *C. americana* (‘Winkler’) | 327.7 | 48.4% | 23,331 | 96.9 | 96.5 |
| *C. americana* (‘Rush’) | 350.2 | 49.0% | 24,562 | 99.0 | 99.0 |
| *C. mandshurica* | 367.7 | 68.7% | 25,923 | 97.1 | 92.2 |
| *C. heterophylla* (Beijing) | 361.9 | 56.7% | 27,591 | 93.5 | 92.0 |
| *C. heterophylla* (Jilin) | 342.9 | 58.0% | 22,319 | 94.7 | 92.7 |
| *C. avellana* (‘TGdL’) | 373.1 | 41.5% | 27,791 | 96.2 | 92.0 |
| *C. avellana* (‘Tombul’) | 369.8 | 38.1% | 27,270 | 97.0 | 76.8 |

The total number of predicted genes are within the range previously reported in *Corylus*. In addition, the BUSCO scores for ‘Rush’ and ‘Winkler’ are substantially higher than the scores reported for any previous *Corylus* genome annotation, and are very close to the assembly BUSCO scores. This suggests not only that these annotations are highly complete, but also provides support for the accuracy of the predictive procedure described above.

### Comparison across the Corylus genus

Comparisons across multiple species can be made simultaneously, or in a pairwise manner. Using the five previously published *Corylus* assemblies, which contain comparable annotations to the genome presented here, together with the closely related species *Betula pendula* and outgroup *Malus domestica*, the GENESPACE package (Lovell et al., 2022) was utilized to visualize syntenic relationships in both manners, as well as construct a pangenome assembly anchored to *C. americana* ‘Winkler’. Fig. 2 illustrates the general result that across these *Corylus* genomes, there is almost universal synteny between all defined blocks, illustrating the high degree of relatedness between each pair of species. Differences in pseudomolecule numbering is most likely an artifact of chromosome numbering typically being made according to descending physical contig size. *Corylus* chromosome sizes are generally very similar, and assemblies had slight variations in contig lengths. Fig. 3 shows dot plots representing each pairwise comparisons between syntenic blocks for each of the seven *Corylus* genomes. Despite the high degree of synteny across all *Corylus* species on a chromosome scale, it is clear that *C. americana* exhibits nearly perfect synteny with the two *C. avellana* assemblies. This decreases marginally when compared against the two *C. heterophylla* assemblies, with small inversions evident on a number of chromosomes. *C. mandshurica* very clearly exhibits the least amount of synteny, with large inversions and rearrangements of syntenic blocks across most chromosomes.

**Figure 2.**
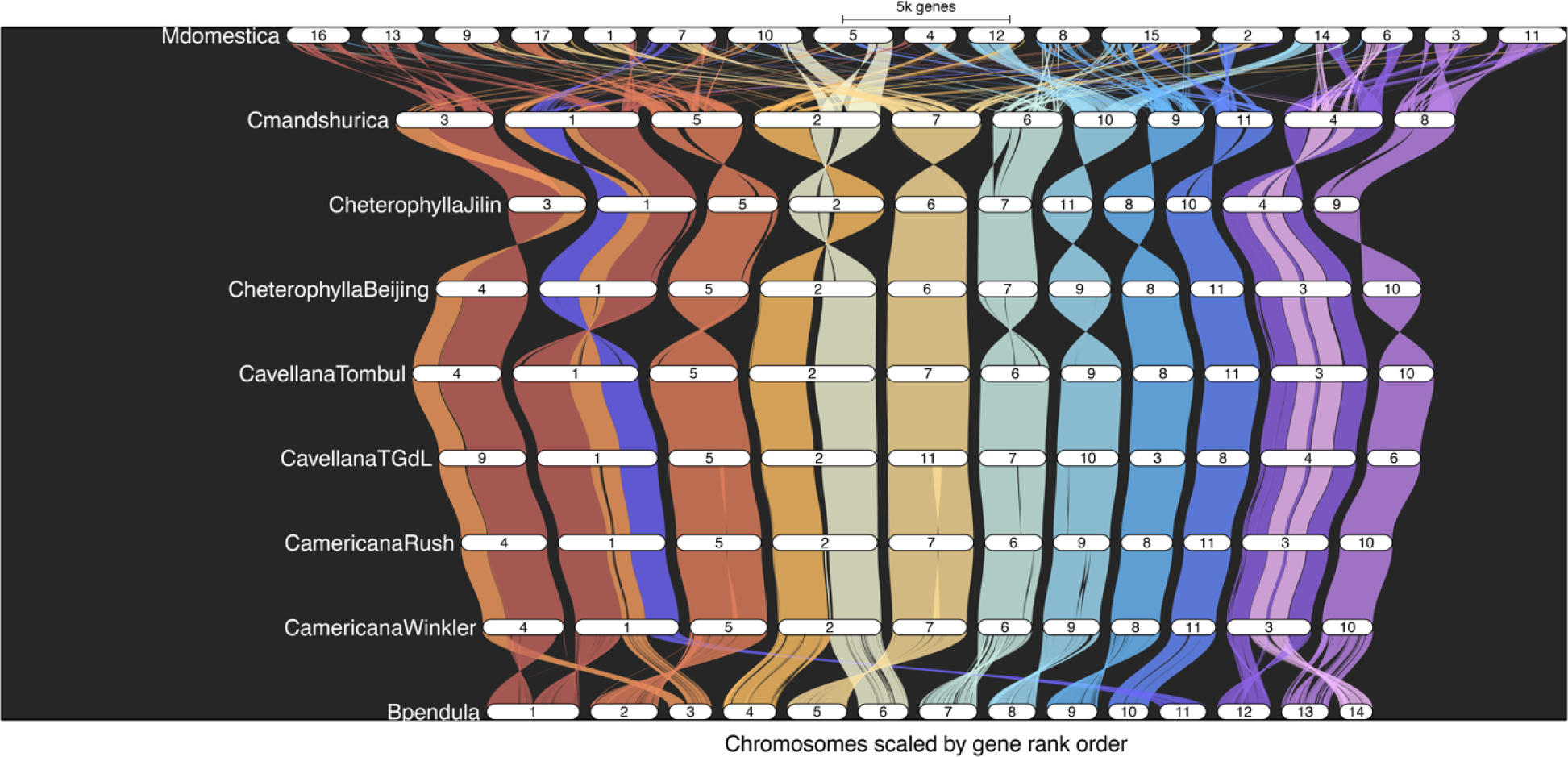
Riparian plot generated by GENESPACE. Single copy orthologs were used to generate and visually compare synteny between ‘Rush’ and ‘Winkler’, the five other published *Corylus* genomes, *Betula pendula* (a close relative in the *Betulaceae*), and the outgroup *Malus domestica*. In addition to illustrating consistent genome-wide inter-species synteny across the 11 chromosomes of *Corylus*, this plot clearly illustrates the large-scale relationships between these 11 chromosomes, and the 14 chromosomes of *B. pendula*.

**Figure 3.**
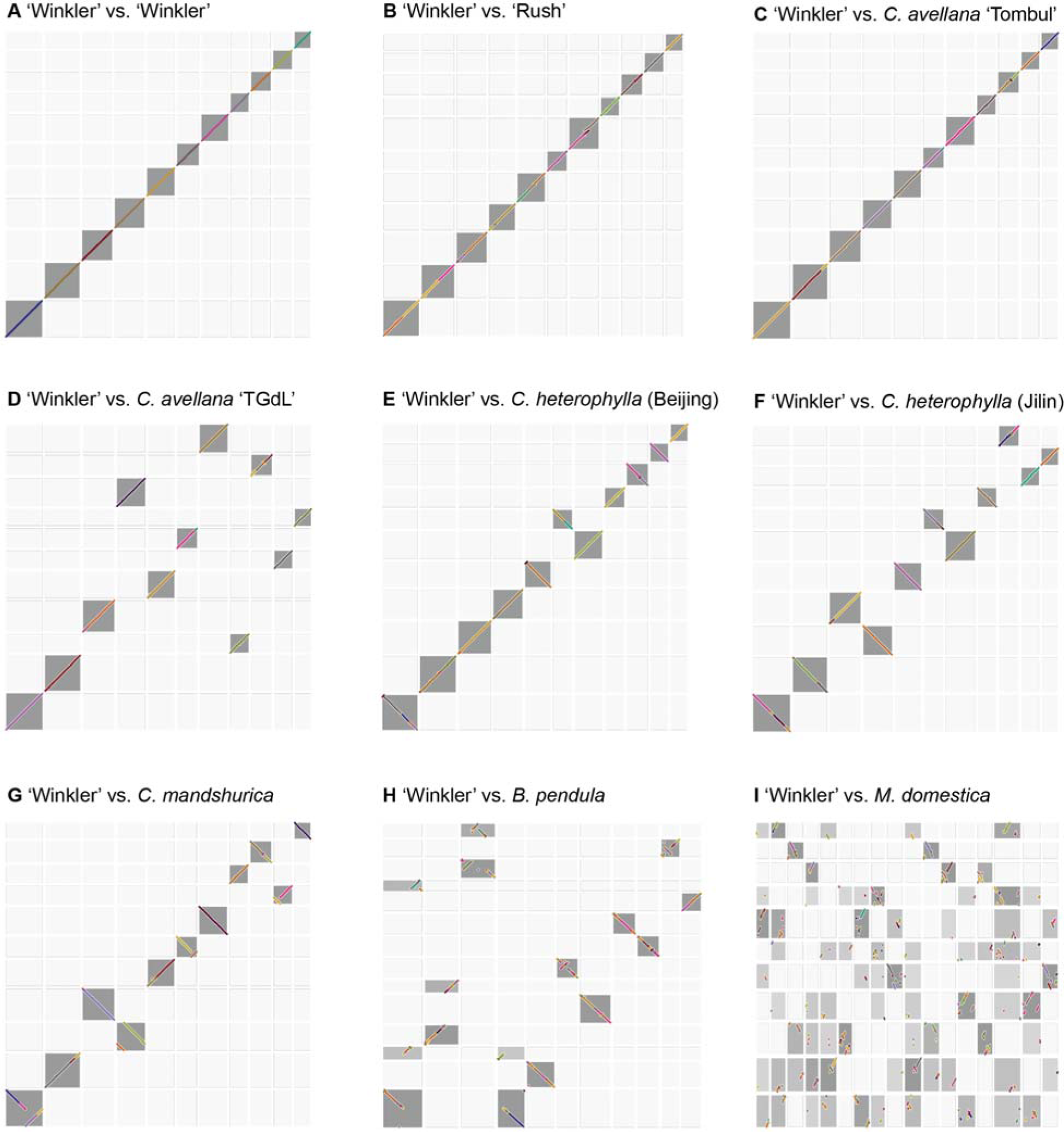
Pairwise dotplots visualizing pairwise BLAST hits between C americana ‘Winkler’, and each of the other eight genomes included in the GENESPACE analysis. Target and query genes which were identified as belonging to the same orthogroup are plotted, and color coded by sequence similarity. **A:** ‘Winkler’ vs. ‘Winkler’; **B:** ‘Winkler’ vs. ‘Rush’; **C:** ‘Winkler’ vs. *C. avellana* ‘Tombul’; **D:** ‘Winkler’ vs. *C. avellana* Tonda Gentille delle Longhe; **E:** ‘Winkler’ vs. C. heterophylla (Beijing); **F:** ‘Winkler’ vs. *C. heterophylla* (Jilin); **G:** ‘Winkler’ vs. *C. mandshurica*; **H:** ‘Winkler’ vs. *B. pendula*; **I:** ‘Winkler’ vs. *M. domestica*.

With respect to *B. pendula*, this figure makes clear the specific large-scale rearrangements which relate the 14 chromosomes of *B. pendula* to the 11 of *Corylus* spp. Specifically, chromosomes 1 and 3 in *B. pendula* were split to form chromosomes 1 and 4 in *Corylus* spp., with *B. pendula* chromosome 11 also being fused with the end of chromosome 1 in *Corylus*. In addition, chromosomes 4 and 6 in *B. pendula* together constitute chromosome 2 in *Corylus*, while chromosomes 12 and 14 constitute chromosome 3 in *Corylus*.

### *Single copy orthologs unique to* C. americana

It is also possible to use the orthogroups generated by OrthoFinder to determine, instead of synteny, specifically those putative genes in *C. americana* which possess no ortholog in any currently annotated *Corylus* species, nor the other two outgroups included in this analysis: *B. pendula* and *M. domestica*. This subset of the transcriptome consists of 66 predicted single copy orthologs present in both ‘Rush’ and ‘Winkler’, but absent in all other analyzed genomes (Supplementary File 1). By filtering for only those single copy orthologs present in the predicted gene sets for ‘Rush’ and ‘Winkler’ limits the possibility that these unique genes are an artifact of spurious assembly errors in either of the independent ‘Rush’ or ‘Winkler’ assemblies. Similarly, including a diversity of assemblies of other *Corylus* species, as well as the *Betula* and *Malus* outgroups, limits the potential that these genes have been identified as unique simply as a result of more successful gene prediction for ‘Rush’ and ‘Winkler’, which due to the high BUSCO scores for the annotations reported here, would otherwise be a concern.

Many of these genes logically did not return hits when the Viridiplantae database was queried using NCBI-BLAST. Several, however, are predicted to be involved in defense response pathways. Of particular note is the gene CamerWinkler.08G009300 (homologous to CamerRush.08G012000.1 in ‘Rush’), which is characterized as involved in “defense response to fungus”. Given *C. americana*’s high level of resistance to the endemic fungal pathogen EFB, this gene would be a valuable target for future functional characterization.

### *Contributions of* C. americana *and* C. avellana *to current Midwest germplasm*

1,343 individual plants were sequenced using GBS, and SNP markers were called using TASSEL. Sampled plants represented wild *C. americana*, cultivars from breeding programs in Oregon, New Jersey, and Minnesota, and F_1_ populations between these cultivars. A biplot for the first two principal components of the distance matrix constructed using these markers is shown in Fig. 4. This visualization makes evident the wide genetic diversity represented in current Midwestern hazelnut varieties, and lends support to the significant *C. avellana* contribution to specific selections, such as Rose9-2. Many currently named Midwestern varieties, on the other hand, cluster extremely closely with ‘Rush’, ‘Winkler’, and wild *C. americana* from the DNR, suggesting that while an interspecific cross may have occurred in the past, subsequent selection of progeny greatly favored *C. americana* genetic contributions.

**Figure 4.**
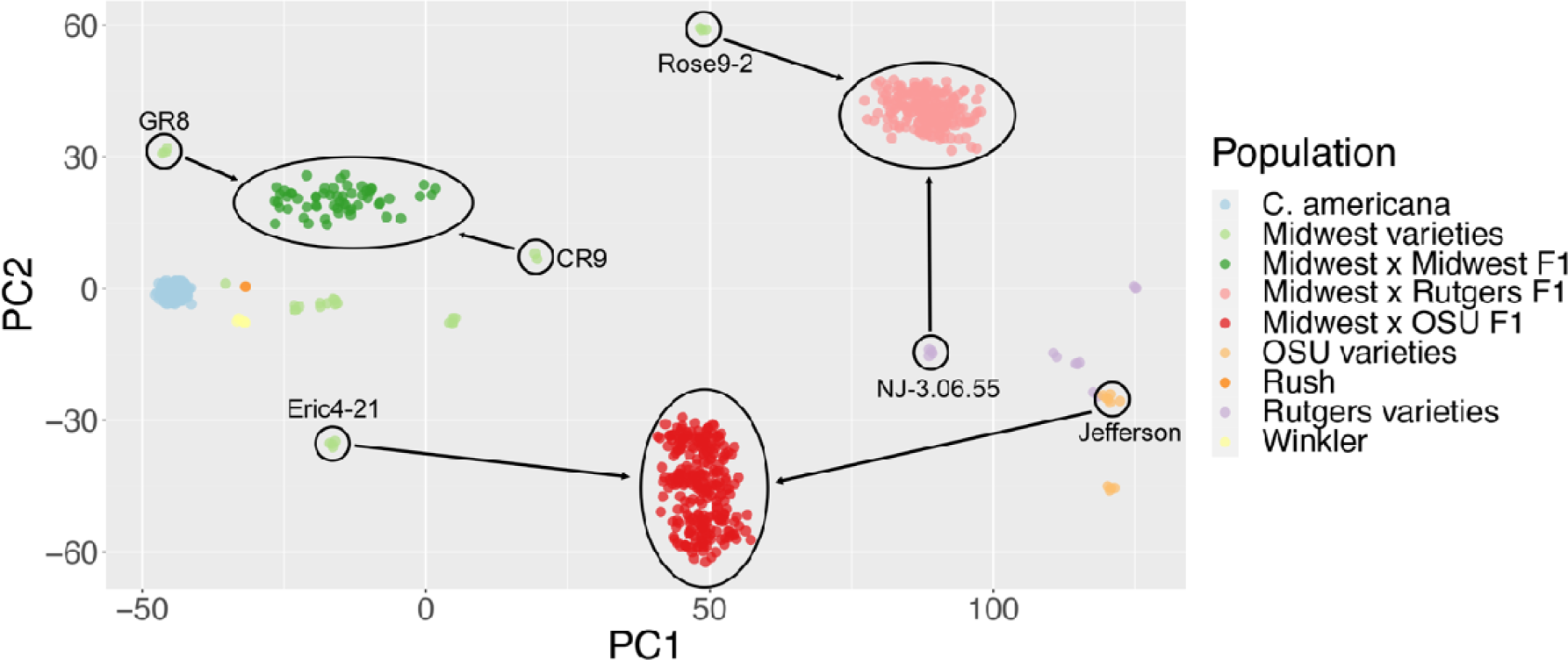
PCA biplot of SNPs identified in *C. americana, C. avellana* and interspecific hybrids. On the left, wild *C. americana* sourced from the DNR clusters together with ‘Rush’ and ‘Winkler’, along with a number of Midwestern varieties and an F_1_ family produced through a cross between two of them (light and dark green dots, respectively). In the middle, an outlier Midwest variety (Rose9-2) exhibits PC1 scores similar to an F_1_ between Eric4-21 (a Midwest variety that appears similar to *C. americana*) x Jefferson (a *C. avellana* variety from Oregon State University). On the right are *C. avellana* varieties from Oregon State University and Rutgers University, along with an interspecific F_1_ population between a Rutgers cultivar and Rose9-2. This plot illustrates that while most Midwestern varieties appear to be genetically quite similar to *C. americana*, there is substantial diversity among them (more so than the limited number of *C. avellana* varieties included in the analysis) with some appearing quite similar to F_1_ interspecific F_1_ hybrids – a finding supported by the historical record of interspecific hybridization in Midwestern breeding efforts.

### Phylogenetic analysis of sporophytic self-incompatibility

The locus regulating sporophytic self-incompatibility (SSI) in hazelnut has been fine-mapped in *C. avellana* (Hill et al., 2021) and *C. heterophylla* x *C. avellana* interspecific hybrids (Hou et al., 2022). These studies identified three MIK2 homologues believed to be responsible for SSI in these two *Corylus* species. In order to examine their phylogenetic relationship, multiple sequence alignment was performed between the top BLAST hit for each of these three genes and all currently available *Corylus* transcriptomes. Outgroups were included as above for *B. pendula* and *B. oleracea*, which both also exhibit SSI (Hynynen et al., 2010; Kitashiba & Nasrallah, 2014), as well as *M. domestica*, a more distantly-related species that is a well-studied example of gametophytic self-incompatibility (Cheng et al., 2006; Minamikawa et al., 2010). This phylogenetic tree is shown in Fig. 5, and illustrates the genus-specific evolution of SSI in *Corylus*, including several clades containing orthologous sequences only found in *Corylus* species.

**Figure 5.**
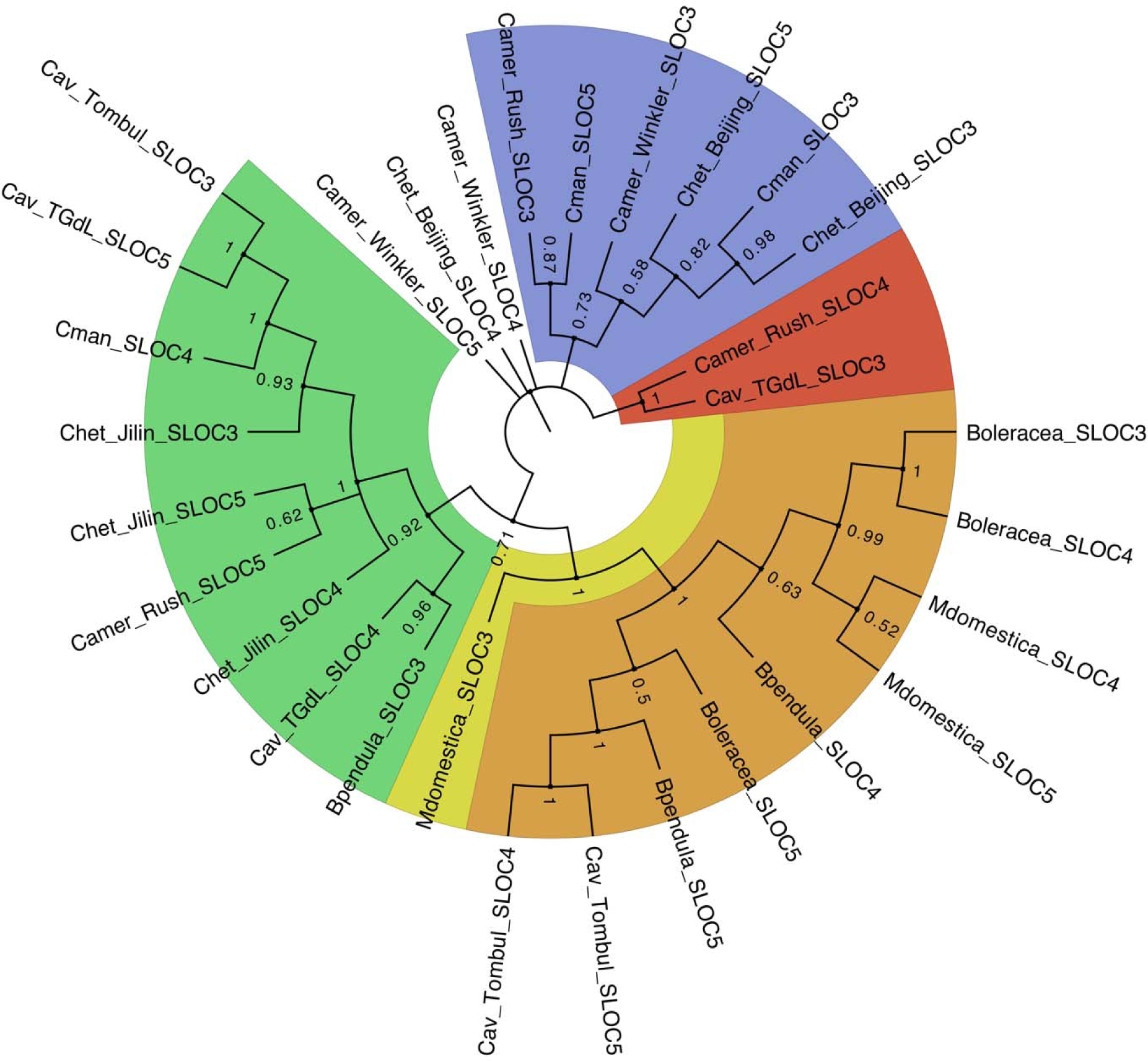
Phylogenetic tree showing the evolution of genes comprising the self-incompatibility locus in *Corylus* across the seven available *Corylus* genomes, with *B. pendula, M. domestica*, and *B. oleracea* included as outgroups. Node labels represent bootstrap values based on 100 permutations. Color overlays illustrate evident clades, and highlight the fact that several lineages of S locus genes are entirely specific to *Corylus*.

## Discussion

This assembly is a critical step in developing the use of next-generation sequencing data in the improvement of *C. americana*, while also enabling comparisons of structural variation across *Corylus* species.

### Variable contiguity of two de novo assemblies

The fragmentation of the ‘Rush’ assembly by the 3D-DNA pipeline (using Arima Hi-C sequence data) was unexpected, given the widespread and successful use of this method in building chromosome-scale assemblies (Ghurye et al., 2019), and in particular, the substantial improvements in contiguity that was observed when using this data to refine the initial ‘Winkler’ assembly. Two possible explanations for this discrepancy suggest themselves. First, the ‘Rush’ accession generated less absolute sequence compared to the ‘Winkler’ assembly on all sequencing platforms, and as a result there was less sequence data to include in each bioinformatic step. The raw assemblies generated by hifiasm were indeed also more contiguous for Winkler compared to ‘Rush’. Nevertheless, with coverage >40x for all sequence data types, the failure of Hi-C sequence to improve contiguity would not appear to be solely a consequence of a lack of data. A second likely compounding factor, however, is a difference in the relative heterozygosity of ‘Rush’ and ‘Winkler’. Heterozygosity is known to complicate phased genome assembly algorithms which use HiFi and Hi-C sequence data (Garg, 2021). Using the 53,696 GBS-derived SNPs described above it was possible to directly estimate genome-wide heterozygosity as a ratio of heterozygous to total polymorphic sites. By this metric, 22% heterozygosity was observed for ‘Rush’, while only 17% of polymorphic sites were heterozygous in ‘Winkler’, suggesting that this could have contributed to the challenges in scaffolding ‘Rush’ using Hi-C data.

Regardless of the specific underlying factors complicating this stage of genome assembly, it is clear that even in diploid species with relatively small genome sizes, integrating chromosomal conformation capture sequencing data into genome assembly pipelines is not a universally applicable method. At the same time, the use of high-coverage long-read sequence data, combined with modern assemblers for scaffolding and polishing chromosome-scale genomes, has substantial capacity to generate high-quality reference assemblies, even in outcrossing, heterozygous organisms.

### PCA-derived population structure

This study reports the first combined analysis of relatedness between commercial varieties of hazelnuts from the three main breeding programs in the United States: Oregon State University, Rutgers University, and the Upper Midwest Hazelnut Development Initiative. This evaluation of population structure, by combining a diversity of clonal varieties with F_1_ full-sib families produced through multiple pairwise controlled crosses of these clones, help elucidate a long outstanding question regarding the degree to which Midwest-bred hazelnut accessions remain closely related to *C. avellana*. 20^th^ century Midwest breeding populations included interspecific crosses with *C. avellana*, but the precise frequency of such hybridization events, and the fact that they were often followed by several generations of open pollination within *C. americana* dominant breeding orchards, has left this question unresolved. Fig. 4 shows clearly that the majority of these Midwest accessions are indeed very closely related to wild *C. americana* sourced from the Department of Natural Resources, as well as ‘Rush’ and ‘Winkler’ themselves. A clear outlier in this respect is the variety Rose9-2, which very closely resembles other F_1_ interspecific crosses.

### Cross-genus level analysis

Transcriptomic analysis of the seven *Corylus* genomes included in this study reveals a consistent distribution of genes across the 11 chromosomes. This analysis considers nearly half of the recognized species in *Corylus* (Thompson et al., 1996). The observed consistency in gene distribution suggests a relatively recent common ancestor of these hazelnut species. This finding is important for understanding the evolutionary relationships between these species and could have implications for hazelnut breeding and genetic improvement programs by providing a foundation for identifying useful traits and genes that are present in all species.

In addition to identifying general synteny on a genome-wide level, these analyses also made possible the identification of those single copy orthologs with no representation in any other Corylus species. In particular, a gene found in both both ‘Rush’ (CamerRush.08G012000.1) and ‘Winkler’ (CamerWinkler.08G009300) was closely associated with defense response to fungus. In particular, it appears to be a defensin-like (DEFL) gene, cysteine-rich antimicrobial proteins (Tesfaye et al., 2013). This locus is therefore a promising target for future functional characterization, and introgression into existing hazelnut cultivars in order to increase their genetic resistance to fungal pathogens.

### Sporophytic self-incompatibility

Attempts at genetically characterizing the SSI S-locus in *Corylus* have included hybridization-based staining of genes identified in *Brassica oleracea* L., which showed irregular hybridization (Hampson et al., 1996). Similarly, gene expression studies identified only 61% homology between *B. oleracea* alleles and S alleles in *Corylus* (Torello Marinoni et al., 2009). Finally, transcriptomic analysis of the fine-mapped S-locus region in multiple *Corylus* species suggest *Corylus* may harbor a novel SSI molecular mechanism that differs from *Brassica* (Hill et al., 2021; Hou et al., 2022). The analysis presented here provides additional evidence for independent evolution of a sporophytic self-incompatibility mechanism that is unique to *Corylus,* and not shared by other members of the *Betulaceae*, nor model species of SSI such as *Brassica oleracea*.

### Conclusion

High-quality, annotated, chromosome-scale genome assemblies are essential tools for utilizing modern genetic methods in the investigation of the molecular control of important traits, as well as the application of advanced breeding methods. These two assemblies provide an important resource which we hope will enable the application of such methods to the study and improvement of both *C. americana*, as well as interspecific hybrids which expand the range over which hazelnuts can be cultivated. Our analysis demonstrates highly conserved genome-wide synteny across *Corylus* species. As such, genomic tools, analyses and resources developed in one species may be more broadly useful to breeding programs in other species within the genus.

## Experimental procedures

### Plant material collection

Tissue was collected from *C. americana* accessions ‘Rush’ and ‘Winkler’ maintained at the National Clonal Plant Germplasm Repository (NCGR) in Corvallis, Oregon (PI 557022 and PI 557019, respectively). ‘Rush’ is a specimen collected around 1900 by J.F. Jones in Lancaster, Pennsylvania, while ‘Winkler’ was collected in 1910 by Wendell Williams in Danville, Iowa (Molnar, 2011). These two varieties were historically significant in early- and mid-20^th^ century hazelnut breeding programs, and as such represent genetically relevant, pure *C. americana* selections, which also reflect the wide geographic distribution of *C. americana* in the Eastern U.S. Today, ‘Rush’ remains a widely used source of EFB resistance (Bhattarai et al., 2017). Photos of the clones located at the NCGR are shown in Fig. 1A. Tissue was collected on April 13^th^, 2020, from bushes that had been etiolated for three days prior to sampling in order to minimize the concentration of volatiles and secondary metabolites in leaf tissue. Samples were immediately flash-frozen using liquid nitrogen and stored at -80°C until DNA extraction.

### Long-read sequencing

High-molecular weight DNA was extracted following the protocol described by Vaillancourt & Buell (2019). In brief, leaf tissue was homogenized in liquid nitrogen, lysed with Carlson lysis buffer, and purified with chloroform and Qiagen Genomic-tips (Qiagen N.V., Venlo, The Netherlands). Purity of extracted DNA was assessed spectrophotometrically using a NanoDrop™ One (ThermoFischer Scientific, Waltham, Massachusetts). DNA was quantified a using Qubit™ dsDNA High Sensitivity kit (ThermoFisher Scientific, Waltham, Massachusetts), and diluted and assessed for size using an Agilent FemtoPulse System (Santa Clara, California).

Pacific Biosciences HiFi libraries were prepared according to PN 101-853-100 Version 03 (Pacific Biosciences, Menlo Park, California). Modifications included shearing with a Covaris g-TUBE (Covaris, Woburn, Massachusetts) and size selecting with BluePippin (Sage Science, Beverly, MA). Libraries were sequenced on a PacBio Sequel II using the Sequel Polymerase Binding Kit 2.2 at the University of Wisconsin-Madison Biotechnology Center’s DNA Sequencing Facility. Oxford Nanopore libraries were prepared following the Native Barcoding Expansion protocol (Oxford Nanopore Technologies, Oxford, United Kingdom). The library was sequenced on a R9.4.1 flowcell on an Oxford Nanopore PromethION, also at the UW-Madison Biotechnology Center. Partway through the sequencing run, DNA was flushed with the Oxford Nanopore Technologies Flow Cell Wash Kit (EXP-WSH004), and additional library was loaded.

### Hi-C sequencing

Nuclei were extracted from flash-frozen leaf tissue using a Sigma CelLytic™ PN Plant Nuclei Isolation/Extraction Kit (Sigma-Aldrich, Burlington, Massachusetts). Crosslinking was performed following low-input protocols from Arima Genomics (Arima Genomics, Carlsbad, California). Crosslinked nuclei were quantified using a Qubit™ dsDNA High Sensitivity kit, samples were sheared to 600bp, and library preparation was performed using a KAPA® Hyper Prep kit (Roche, Basel, Switzerland). The final library was assessed for quality using the Agilent TapeStation System with a D1000 kit (Agilent Technologies, Santa Clara, California), and paired-end, 150 bp paired-end sequences were generated using an Illumina NovaSeq 6000 (Illumina, San Diego, California).

### RNA extraction, library preparation and sequencing

Prior to RNA extraction, Tissuelyser II adapters, safelock tubes, and 5mm stainless steel beads were chilled at -80°C for 2 hours. Working on dry ice, leaf and kernel tissue was added to prechilled tubes and disrupted using the Tissuelyser II (Qiagen N.V., Venlo, The Netherlands) for 2 rounds at 30Hz for 1 minute each round. Samples were then processed using the Qiagen Plant RNeasy (Qiagen N.V., Venlo, The Netherlands) with an on-column DNA digest. Total RNA was assayed for purity and integrity using a NanoDrop One™ Spectrophotometer and Agilent 2100 Bioanalyzer (Agilent Technologies, Santa Clara, California), respectively. RNA libraries were prepared from samples that met the Illumina TruSeq® Stranded Total RNA Sample Preparation Guide (15031048 E) input guidelines using the Illumina TruSeq® Stranded Total (Plant) RNA Sample Preparation kit (Illumina Inc., San Diego, California, USA). For each library preparation, cytoplasmic, mitochondrial and chloroplast ribosomal RNA was removed using biotinylated target-specific oligos combined with paramagnetic beads tagged with streptavidin. Following purification, the reduced RNA was fragmented using divalent cations under elevated temperature. Fragmented RNA was copied into first stranded cDNA using SuperScript II Reverse Transcriptase (Invitrogen, Carlsbad, California, USA) and random primers. Second strand cDNA was synthesized using a modified dNTP mix (dTTP replaced with dUTP), DNA Polymerase I, and RNase H. Double-stranded cDNA was cleaned with AMPure XP Beads (1X) (Beckman Coulter, Brea, California,). The cDNA products were incubated with Klenow DNA Polymerase to add a single “A” nucleotide to the 3’ end of the blunt DNA fragments. Unique dual indexes (UDI) were ligated to the DNA fragments and cleaned with two rounds of AMPure XP beads (0.8X). Adapter-ligated DNA was amplified by PCR and cleaned with AMPure XP beads (0.8X). Final libraries were assessed for size and quantity using an Agilent DNA1000 chip and a Qubit® dsDNA HS Assay Kit (Invitrogen, Carlsbad, California, USA), respectively. Libraries were standardized to 2 nM, and 150-bp paired-end sequencing was performed on an Illumina NovaSeq 6000.

### Genome assembly

An initial assembly was created using PacBio CCS reads with the program hifiasm v0.16.0-r369 (Cheng et al., 2021) on a Unix server with 160 cores and 3TB of RAM. This draft assembly exhibited higher contiguity than references generated with Canu using ONT reads, HiCanu, and Flye using either HiFi or ONT reads. The program purge_haplotigs v1.1.1 (Roach et al., 2018) was then utilized to reduce artificial genome duplication caused by incorporation of non-collapsed haplotigs in the final assembly. Hi-C reads were aligned, filtered, and binned using the haplotig-purged assembly using the program juicer (Durand et al., 2016). Contigs were then scaffolded and ordered using the program 3D-DNA (Dudchenko et al., 2017).

While integration of Hi-C data led to improved contiguity and reduced artificial duplication in the ‘Winkler’ assembly, contigs in ‘Rush’ were fragmented following the use of juicer and 3D-DNA. As a result, this step was omitted for ‘Rush’. The assemblies were next iteratively polished three times with Racon (Vaser et al., 2017), using the original PacBio reads. We then used a recently constructed linkage map to detect erroneous inversions in the physical assemblies (Brainard et al., 2023), using visual inspection of heatmaps of recombination frequencies in an F_1_ population. The program Ragtag v2.1.0 (Alonge et al., 2022) was then used to correct observed errors using ONT reads which overlapped any identified inversions. Initial quality assessments were made using BUSCO v5.4.3 (Manni et al., 2021) with the eudicots_odb10 database, and quast v5.2.0 (Mikheenko et al., 2018) with default parameters.

### Genome annotation

Transcript assemblies for both ‘Rush’ and ‘Winkler’ were made from ∼200M 2 × 150 bp stranded paired-end Illumina RNAseq reads using the software PERTRAN, which conducts genome-guided transcriptome short read assembly via GSNAP (Wu & Nacu, 2010) and builds splice alignment graphs after alignment validation, realignment, and correction. These outputs were subsequently used to construct 37,794 and 36,419 transcript assemblies for ‘Rush’ and ‘Winkler’, respectively, using the program PASA (Haas et al., 2003). Loci were determined by transcript assembly alignments and/or EXONERATE alignments of proteins from *Mimulus guttatus, Arabidopsis thaliana, Gossypium raimondii, Betula platyphylla, Carya illinoinensis, Castanea dentata, Quercus rubra, Prunus persica, Fragaria vesca, Glycine max, Medicago truncatula, Vitis vinifera, Liriodendron tulipifera, Juglans microcarpa, Populus trichocarpa, Beta vulgaris, Solanum lycopersicum, Sorghum bicolor, Oryza sativa*, and Swiss-Prot (release 2022_04 of eukaryote proteomes), with up to 2k bp extension on both ends unless extending into another locus on the same strand. Alignments were made to the respective genome assemblies following softmasking for repetitive elements. The repeat library consisted of *de novo* repeats identified by RepeatModeler2 (Flynn et al., 2020), generated using both the ‘Rush’ and ‘Winkler’ assemblies, as well as *C. americana* repeats identified in the databases RepBase and Dfam. Gene models were predicted by homology-based predictors: FGENESH+ (Salamov & Solovyev, 2000), FGENESH_EST (which is similar to FGENESH+, but uses EST to compute splice site and intron input instead of protein/translated ORF), EXONERATE (Slater & Birney, 2005), PASA assembly ORFs (a homology constrained ORF finder), and AUGUSTUS (Stanke et al., 2006) trained by the high confidence PASA assembly ORFs, and with intron hints from short read alignments. The best-scored predictions for each locus were selected using multiple positive factors including EST and protein support, and overlap with repeats as a negative factor. The selected gene predictions were improved by PASA. The improvement included adding UTRs, splicing correction, and adding alternative transcripts. PASA-improved gene model proteins were subject to protein homology analysis to the above-mentioned proteomes to obtain a Cscore and protein coverage. Cscore is a protein BLASTP score ratio to the mutual best hit (MBH) BLASTP score, and protein coverage is the highest percentage of protein aligned to the best homologs. PASA-improved transcripts were selected based on Cscore, protein coverage, EST coverage, and their CDS overlap with repeats. The transcripts were selected if their Cscore and protein coverage were ≥ 0.5 or if covered by ESTs. For gene models whose CDS were overlapped by repeats by more than 20%, their Cscore was required to be at least 0.9, and homology coverage at least 70%, in order to be selected. The selected gene models were subject to Pfam analysis, and gene models whose proteins were more than 30% overlapped by Pfam TE domains were removed as weak gene models. Incomplete gene models, low homology supported gene models without full transcriptome support, short single exon (< 300 bp CDS) with neither protein domains nor good expression, and repetitive gene models without strong homology support were manually filtered out.

### Functional annotation

Every peptide sequence in the dataset was analyzed with a computational pipeline that includes the standard InterProScan (Jones et al., 2014) suite of programs to determine protein domains and other sequence features, E2P2 (Chae et al., 2014; Schläpfer et al., 2017) for enzyme assignments (EC), and PathoLogic (Karp et al., 2021) for metabolic pathway assignments. Additional processing was used to determine Eukaryotic Orthologous Groups (KOG) gene assignment using a modified mutual best hit algorithm. Results of the InterProScan calculations were used to assign standard InterPro protein domain associations and from these, gene ontology (GO) terms. Protein domains inferred from these calculations were used to develop a putative gene functional assignment which includes a count of the multiplicity of the assignment in the proteome set.

### Genotyping by sequencing

To better compare genomes of *C. avellana* and *C. americana*, we used GBS to genotype a population of interspecific hybrids and a population of *C. americana.* Tissue from 1,343 samples of breeding lines from the Upper Midwest Hazelnut Development Initiative (UMHDI), Oregon State University, in Corvallis, Oregon, and Rutgers University, in New Brunswick, New Jersey, along with full-sib F_1_ populations derived from controlled crosses between these varieties, and a wild Midwestern population of *C. americana* sourced from the Wisconsin Department of Natural Resources and planted in Barneveld, WI, was sampled following budbreak in May of 2020. Genomic DNA was extracted and libraries for genotyping-by-sequencing were prepared using a double digestion with the restriction enzymes *Nsi*I and *Bfa*I following the methodology described by Elshire et al. (2011). This specific double digest was selected based on analysis of k-mer distributions of sequence libraries generated with *ApeK*I alone, *Nsi*I and MspI, *Pst*I and *Bfa*I, *Pst*I and *Msp*I, *ApeK*I and *Bfa*I, and *ApeK*I and *Msp*I. Illumina GBS barcodes and adapters were ligated, and paired-end reads (2 × 150Lbp, 10 million reads / sample) were generated using an Illumina NovaSeq 6000.

Due to the high levels of synteny across the genomes described above, direct use of Illumina sequence data to infer variable degrees of interspecific hybridization, using tools such as sppIDer (Langdon et al., 2018), is limited. Biallelic SNPs were therefore identified using the TASSEL GBSv2 pipeline (Bradbury et al., 2007) and filtered for missing data (< 10% across all samples), minor allele frequency (> 0.05), linkage disequilibrium (r^2^ < 0.75), and allele depth (80% of samples with a depth > 8) using bcftools (Danecek et al., 2021). Population structure analysis using principal components analysis (Price et al., 2006) has been shown to be a simple and efficient alternative to more complex model-based approaches such as STRUCTURE (Falush et al., 2007) and ADMIXTURE (Alexander et al., 2009). This filtered VCF file was therefore converted to a Euclidean distance matrix using R (R Core Team, 2021), and multi-dimensional scaling (which permits some missing data) was then performed to obtain measures analogous to scores along the first two principal components of the distance matrix.

### Comparative analyses of Corylus

Comparisons across the *Corylus* genus were made utilizing the five currently available chromosome-scale genome assemblies, as well as the *C. americana* assemblies reported here for ‘Rush’ and ‘Winkler’. This included two *C. avellana* assemblies (cultivars ‘Tombul’ (Lucas et al., 2021) and ‘Tonda Gentile della Langhe’ (Pavese et al., 2021)), two *C. heterophylla* assemblies (an accession from Siping City, Jilin (Liu et al., 2021) and an accession from Yanqing, Beijing (Zhao et al., 2021)), and one *C. mandshurica* assembly (a wild specimen from Xinglong (Li et al., 2021)). All of these assemblies were annotated using similar *ab initio* prediction methods which combined RNAseq data and protein homology data, and thus their transcriptomes were well-suited to comparative analyses with ‘Rush’ and ‘Winkler’.

GENESPACE v0.94 (Lovell et al., 2022) was used to investigate genome-wide syntenic relationships, with *Betula pendula* (silver birch) and *Malus domestica* (apple) included as outgroups (gene models being obtained from https://genomevolution.org/coge/ and https://www.rosaceae.org, respectively). In brief, GENESPACE utilizes the program OrthoFinder (Emms & Kelly, 2019) to identify orthogroups from predicted gene models, and then parses orthologs to define syntenic blocks across species using BLAST and MCScanX (Wang et al., 2012). Genes in *C. americana* with no orthologous sequences in any other *Corylus* species were also identified using OrthoFinder.

### Phylogenetic analysis of sporophytic self-incompatibility

Sequences for three S-locus genes (MIK3 homologues) reported by Hou et al. (2022) were obtained from the transcriptome reported in Zhao et al. (2021). NCBI BLAST v2.6.0+ (Altschul et al., 1990) was used to identify homologous sequences in all *Corylus* species, as well as *B. pendula, M. domestica*, and *Brassica oleracea*. These latter three species were included as variably-related outgroups which also exhibit self-incompatibly: SSI in the case of *B. oleracea* and *B. pendula*, and GSI in the case of *M. domestica*. The *B. oleracea* transcriptome was obtained from http://brassicagenome.net/. Multiple sequence alignments were made using muscle v5.1 (Edgar, 2021). These 84 alignments were then imported into MEGA11 (Tamura et al., 2021) and used to build a phylogenetic tree using maximum likelihood and the Jones-Taylor-Thornton matrix-based model (Jones et al., 1992). A bootstrap consensus tree was inferred from 100 replicates, in which branches present in less than 50% of these replicates were collapsed. Trees were visualized and clades labeled using the program FigTree v1.4.4.

## Acknowledgments

We thank Al Kovalevski, Samridhi Chaturvedi and Ashely Yow for providing helpful advice in the performance of phylogenetic analysis. In addition, Nahla Bassil, Shawn Mehlenbacher, Tom Molnar, Malachi Persche, Lois Braun, and Mark Hamann assisted with the collection of tissue samples used in this study. We thank Thomas Hickey and Marissa Nix as well as the staff of the University of Wisconsin–Madison Biotechnology Center DNA Sequencing Facility and the USDA National Clonal Germplasm Repository in Corvallis, OR for logistical support of the research. We thank Chuck and Gerta Zinda for allowing us to study the *C. americana* planted on their property in Barneveld, WI for our research.

## Funding

Sequencing and bioinformatics was supported by the Jewett Prize of the Arnold Arboretum and USDA-NIFA SCRI Grant No. H007913501, with matching funds from the Savanna Institute and The Grantham Foundation for the Protection of the Environment. Genome annotation conducted at the U.S. Department of Energy Joint Genome Institute (https://ror.org/04xm1d337) was supported by the Office of Science of the U.S. Department of Energy under Contract No. DE-AC02-05CH11231.

## Conflict of interest

All authors declare no conflict of interest with the research and findings reported here.

## Author contributions

SHB and JCD designed the analysis. DMS and SHB carried out genome assembly and SNP calling. SHB, TB and SS performed genome annotation. SHB performative comparative genetic analyses and prepared the manuscript, with contributions from all authors.

## Data availability statement

Genome assemblies, as well as short- and long-read sequence data (from Arima Hi-C, Oxford Nanopore, PacBio, and RNAseq libraries) are available through the NCBI Genome & Sequence Read Archive, respectively, under BioProject ID: PRJNA939214 BioSample accessions: SAMN33458638 (‘Rush’), SAMN33458639 (‘Winkler’). Genome assemblies, GFF3 models of predicted genes, functionally annotated peptide sequences, anchored pangenome, and VCF of GBS-derived SNPs are also available at https://doi.org/10.5281/zenodo.7439335. Genomes and annotations are also available through Phytozome: https://phytozome.jgi.doe.gov/info/Camericanavar_winkler_v1_1 (for Winkler); and https://phytozome.jgi.doe.gov/info/Camericanavar_rush_v1_1 (for Rush). Bioinformatic pipelines for genome assembly and polymorphism identification are published at: https://github.com/shbrainard/Camericana_pipelines.

## Notes

https://zenodo.org/deposit/7439335

### Competing Interest Statement

The authors have declared no competing interest.

### Summary of Updates

Updated the analysis of unique single-copy orthologs to include only those overlapping in both 'Rush' and 'Winkler'.

